# Transcriptomic profiling of Parkinson’s disease patient-derived midbrain organoids reveals selective vulnerability of ALDH1A1+ dopamine neurons

**DOI:** 10.64898/2026.11.23.768594

**Authors:** Ghislaine Deyab, Paula Lepine, Taylor M. Goldsmith, Joshua Gertsvolf, Julien Sirois, Carol X.-Q. Chen, Narges Abdian, Jo Anne Stratton, Thomas M. Durcan, Rhalena A. Thomas, Edward A. Fon

## Abstract

Parkinson’s disease (PD) is a neurodegenerative disorder that affects millions worldwide and is broadly characterized by the extensive loss of dopamine neurons (DNs) in the substantia nigra pars compacta (SNpc) relative to neighboring DNs of the ventral tegmental area (VTA). Differences between SNpc and VTA DNs relating to selective vulnerability remains understudied due to limited access to human brain tissue prior to neuronal death, and the limited ability of animal models to fully recapitulate disease processes. Here we demonstrate that diverse DN subpopulations develop spontaneously within human induced pluripotent stem cell (iPSC)-derived midbrain organoids (hMOs). Furthermore, both genetic and sporadic PD patient-derived hMOs exhibit selective loss of *SOX6*+*ALDH1A1*+ but not *CALB1*+ DNs. Surviving *SOX6*+*ALDH1A1*+ DNs in PD hMOs exhibit an increase in mitochondria related gene expression, indicating potential mitochondrial dysfunction, which is not observed in *CALB1*+ DNs. These findings highlight the ability of hMOs to recapitulate key features of disease pathogenesis and identify pathways associated with disease progression and cellular vulnerability prior to cell death in a patient-derived model, something which remains challenging in human postmortem brain tissue.

## Introduction

Midbrain dopamine neurons (DNs) arise from floor plate progenitor cells and develop into three general clusters termed the ventral tegmental area (VTA), the substantia nigra pars compacta (SNpc), and the retrorubral area (RA)^[1, 2]^. These areas differ in axonal projection targets and their regulation of cognitive and motor functions ^[3, 4]^. For example, DNs of the SNpc project to the dorsal striatum forming the nigrostriatal pathway, critical for motor control. Meanwhile, neighboring DNs in the VTA and RA project to the ventral striatum and prefrontal cortex forming the mesocorticolimbic pathway regulating reward seeking, motivation, and emotions. Advances in electrophysiology, target lesioning, and single-cell or single-nuclear RNA sequencing (scRNAseq/snRNAseq) have led to the discovery of functional and transcriptomic heterogeneity both between and within these generally defined DN clusters ^[5–9]^. For example, sub-populations of DNs within the VTA have been shown to dictate different aspects of reward seeking behaviour, learning, and aversion ^[10–12]^. Likewise, populations of DNs along the SNpc dorsal-ventral axis have been shown to dictate different aspects of movement initiation, acceleration, and deceleration ^[8, 9, 13, 14]^, while also exhibiting differential susceptibility to neurodegeneration in Parkinson’s disease (PD)^[15, 16]^.

PD is broadly characterized by the extensive loss of DNs in the SNpc relative to neighboring neurons of the VTA^[17]^, however human postmortem studies reveal a greater loss of DNs in the ventral tier of the SNpc compared to the dorsal tier pointing to underlying differences between DNs along its ventral-dorsal axis^[15, 18]^. As PD is a debilitating neurodegenerative disorder affecting millions worldwide, understanding the molecular profiles of vulnerable and preserved DNs are of particular interest for developing loss prevention or replacement therapies. Efforts to understand differences between degenerating and surviving DNs in PD have used scRNAseq/snRNAseq to identify transcriptomic differences between DN subgroups within the midbrain^[15, 16, 19]^. Both mouse and human midbrain studies suggest that DNs co-expressing *ALDH1A1* and *SOX6* are localized to the ventral SNpc, project to the dorsal striatum, and have the largest risk for neurodegeneration in PD^[9, 20]^. Conversely, preserved DNs exhibit higher expression of *CALB1* and *VGLUT2*, and tend to be localized to the VTA and dorsal SNpc^[3, 4, 9, 21]^. However, studies in the human brain are limited due to technical challenges. For example, postmortem tissue analysis captures a single fixed timepoint, typically at end-stage disease, within what is a decades-long progressive process. This makes it difficult to study the early, disease-relevant processes involved in the initiation and spread of PD pathology. Even in the case of postmortem tissue samples, immunofluorescence provides only a few markers, and snRNAseq requires enrichment to capture enough DNs for quantitative analysis^[15]^. Moreover, mouse genetic models of PD fail to fully replicate human PD pathology, as they do not exhibit DN degeneration without additional insults such as preformed α-synuclein fibril injections or 6-hydroxydopamine lesions^[22]^. These methods raise concerns about the ability of animal models to capture endogenous PD-related processes and highlight the need for human models.

Induced pluripotent stem cell (iPSC)-derived human midbrain organoids (hMOs) are a promising *in vitro* model, capable of generating all brain cells of neuroectodermal lineage, and spontaneously exhibiting multiple aspects of PD pathology^[23–29]^. In previous work, we demonstrated a significant loss of tyrosine hydroxylase (TH)+ neurons in 5-month-old hMOs derived from a patient harboring a triplication of the *SNCA* gene (*SNCA* Trip), encoding for α-synuclein, compared to a CRISPR-corrected isogenic control (Iso CTL)^[26, 27]^. We also show *SNCA* Trip hMOs accumulate phosphorylated α-synuclein and display aberrations in electrophysiological activity highlighting the ability of hMOs to capture complex PD pathology ^[26, 27]^. Here, using scRNA transcriptomics, we report the molecular profile of DN subpopulations in hMOs. We identify a subset of DNs that is lost, and a second population that is preserved in *SNCA* Trip hMOs. These subsets of cells match the profiles of vulnerable and preserved DN populations observed in mice and human postmortem brains, expressing *ALDH1A1* and *CALB1* respectively. Vulnerable and preserved DN populations, and the loss of *ALDH1A1*+ DNs, is similarly observed in hMOs derived from independent iPSC lines from 3 patients with sporadic PD and 4 control individuals. Transcriptional profiling of the surviving vulnerable *ALDH1A1+* DNs shows upregulation of mitochondrial gene expression suggesting mitochondrial dysfunction as an early indicator of degeneration. These results highlight the ability of hMOs to capture complex diseases processes, such as selective vulnerability of DN subpopulations and mitochondrial dysfunction.

## Results

### *SNCA* Triplication midbrain organoids develop distinct dopamine neuron populations with selective vulnerability

To determine if different subtypes of DNs develop in hMOs, we reanalyzed a previously collected scRNAseq dataset derived from *SNCA* Trip and Iso CTL hMOs^[27]^. The raw sequencing files were reprocessed with a 10X Cell Ranger version updated to include intronic read alignments, and filtration of empty droplets, doublets, and low-quality cells (see methods), resulting in 1284 and 2971 cells from the *SNCA* Trip and Iso CTL hMOs respectively. We merged the two datasets and denoised the expression matrix using the adaptively-thresholding low rank approximation (ALRA) algorithm^[30]^. The samples were then integrated using the Seurat canonical correlation analysis and clusters were annotated using a combination of characteristic cell type marker expression, differential gene expression analysis identifying cluster markers, followed by comparison to pathway and cell type libraries from human postmortem, fetal midbrain, and annotation predictions from hMO reference atlases ^[5, 15, 24, 31]^ (Fig.1A). Notably two distinct DN populations were detected both expressing canonical DN markers (*TH, NR4A2)* with additional expression of *SLC17A6, NEUROD2, CALB2,* and *GAD2* in DN-1 relative to DN-2 (Fig.1B). Other key midbrain cell types were observed that express canonical cell type markers; excitatory neurons (*SLC17A6, NEUROD2, CALB2*), inhibitory neurons (*GAD2, SLC32A1, SOX14*), astrocytes (*AQP4, S100B, SOX9*), and oligodendrocytes (*OLIG1, OLIG2, GALR1*) (Fig.1B). Protein expression of markers defining key cell types were also validated by immunofluorescent staining (Fig.1C). Cell types were further confirmed by gene regulatory network analysis using the single cell regulatory network interface and clustering (SCENIC) workflow, which compares gene regulatory networks inferred to be active within single cells ^[32, 33]^. SCENIC analysis inferred increased activity of cell type specific transcription factors for each cluster, with dopamine specific transcription factors highly active in both DN populations (Supplementary Fig.1A-F). Populations of endothelial cells, ependymal cells, fibroblasts, radial glia, stem cells, and a general neuronal population (expressing a mix of excitatory and inhibitory markers) were also observed (Fig.1A,B). We next performed permutation tests to determine which cell type proportions differ significantly between *SNCA* Trip and Iso CTL hMOs. Remarkably, this comparison revealed a complete loss of the DN-2 cluster in *SNCA* Trip hMOs (FDR = 0.001, Log2FD = -6.05), with no change in DN-1 neurons (Fig.1D,E). The loss of DN-2 neurons is consistent with the previously observed decrease in TH+ DNs using immunofluorescent and flow cytometry analysis in 5-month-old *SNCA* Trip hMOs^[26, 27].^

**Figure 1.**
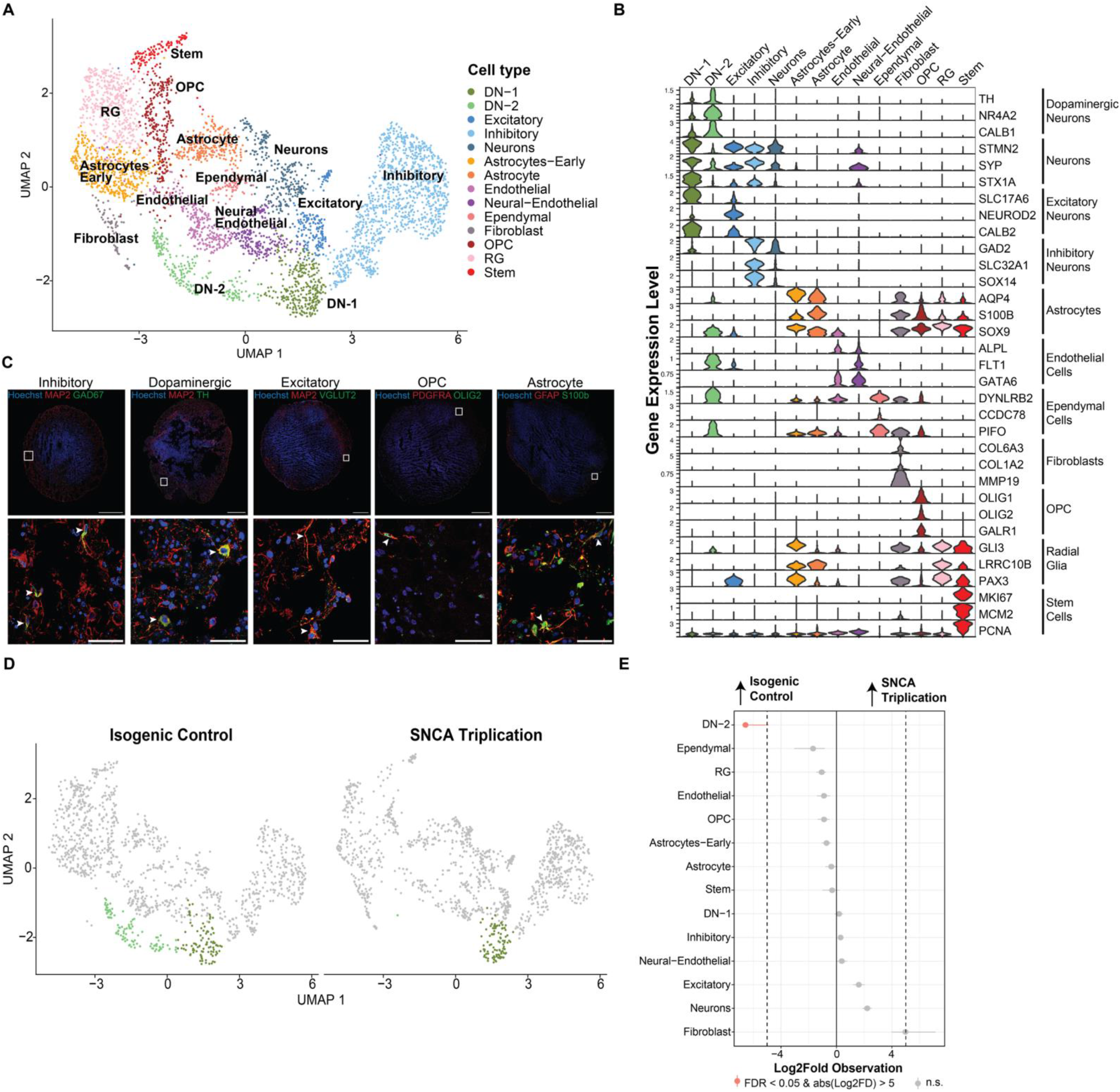
*SNCA* triplication and isogenic control midbrain organoids develop diverse cell types with vulnerable and preserved dopaminergic neuron populations. **(A)** UMAP plot showing annotated cell clusters from *SNCA* Trip and Iso CTL hMOs. **(B)** Violin plot showing expression of canonical cell type markers. **(C)** Immunofluorescent validation of main cell types found via scRNAseq (GAD67+MAP2+ for inhibitory neurons, TH+MAP2+ for dopaminergic neurons, VGLUT2+MAP2+ for excitatory neurons, PDGFRA+OLIG2+ for oligodendrocyte precursor cells, and GFAP+S100b+ for astrocytes). **(D)** Split UMAP showing the loss of DA-2 neurons (light green) in *SNCA* Trip hMOs compared to Iso CTL hMOs. **(E)** Permutation test showing changes in cell proportions between *SNCA* Trip hMOs relative to Iso CTL hMOs. Significant changes in cell proportion are described as those with FDR < 0.05 and log2 fold change > 5.

We next looked at transcriptional differences between DN-1 and DN-2 to further characterize the two populations. For this we isolated both DN-1 and DN-2 cells, rescaled the gene expression, ran a new PCA and UMAP embedding, and reclustered only the two DN populations to examine expression of known broad and subtype specific DN markers. We observe that the DN-2 cluster is enriched in markers previously associated with SNpc DNs and with increased vulnerability in PD animal models including *ALDH1A1, ANXA1,* and *SOX6* ^[3, 5, 6, 34, 35]^ (Fig.2A). Interestingly, DN-1 cells, which are not lost in *SNCA* Trip hMOs, were enriched in markers associated with VTA DNs and decreased vulnerability in PD such as *SLC17A6* and *ADCYAP1* ^[3–5, 9, 21, 35, 36]^ (Fig.2A). We therefore renamed the DN-1 and DN-2 populations as DN-VTA (for VTA-like DNs) and DN-SN (for SN-like DNs) respectively (Fig.2B). These findings indicate that hMOs are capable of developing diverse midbrain DN populations, which differentially express key markers of VTA and SN DNs. Furthermore, the DN-SN population was selectively lost in the *SNCA* Trip hMO ^[3, 4, 6, 9, 15, 16]^.

**Figure 2.**
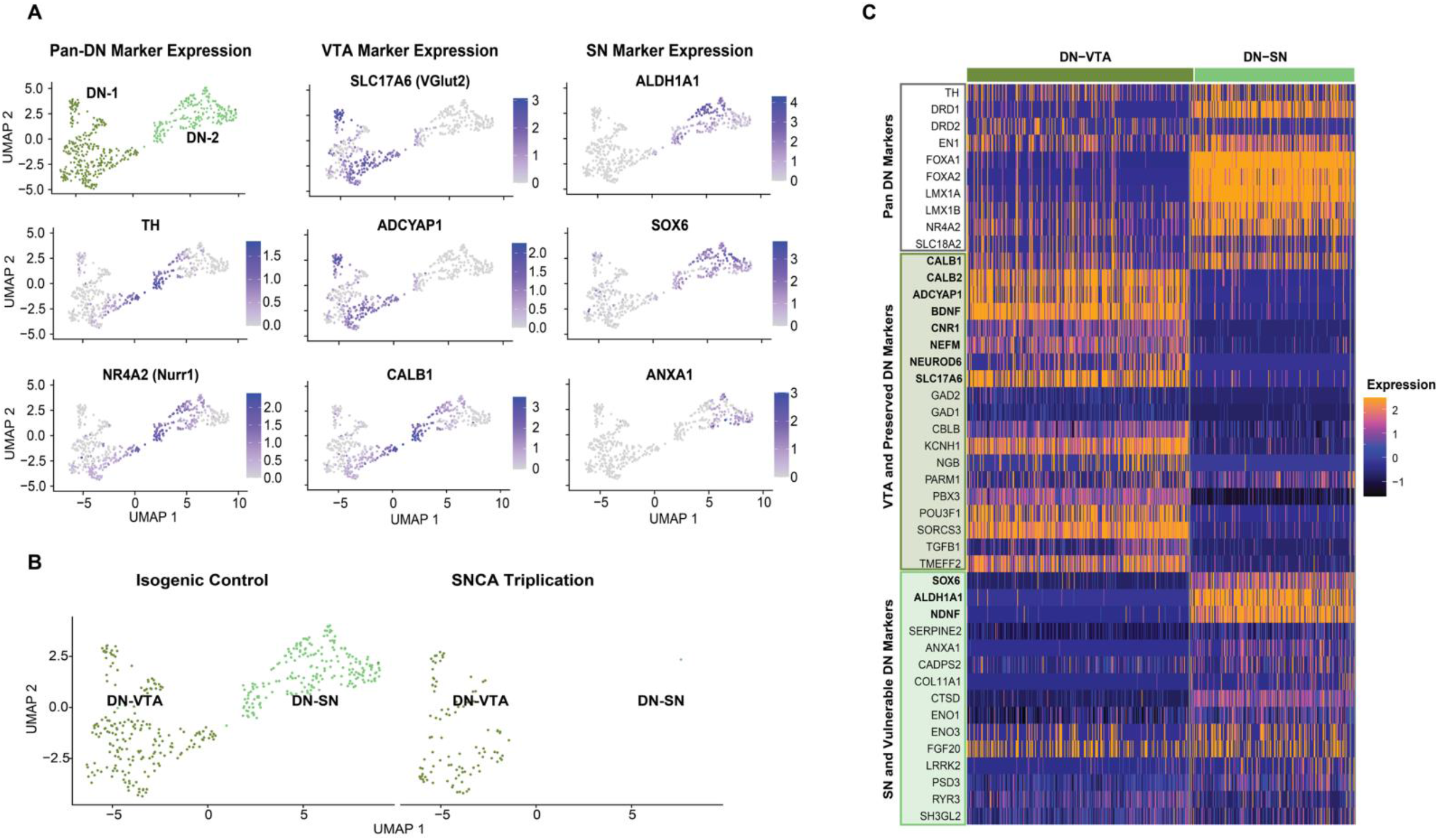
Single cell transcriptomic characterization of dopaminergic neuron subpopulations in *SNCA* triplication midbrain organoids reveals VTA-like and SN-like identities. **(A)** UMAP plot of re-clustered dopaminergic cells followed by expression plots of pan dopaminergic markers, canonical VTA markers, and canonical SN markers across each subtype. **(B)** Split UMAP showing the loss of DN-SNs in *SNCA* Trip hMOs relative to Iso CTL hMOs. **(C)** Heatmap of genes defined in previous studies as either pan dopaminergic (expressed across VTA and SN dopaminergic neurons), highly expressed in VTA and preserved dopaminergic neurons in PD, or highly expressed in SN and susceptible dopaminergic neurons in PD. Bold genes represent those associated with VTA or SN identity, while other genes represent those associated with increased survival or vulnerability in PD, without being a common marker of VTA or SN DNs. **(D)** Volcano plot highlighting DEGs between DN-SN and DN-VTA clusters in Iso CTL hMOs (FDR < 0.001 and log2 fold change > 1.5 or < -1.5).

### Dopamine neuron populations within midbrain organoids resemble mouse and human midbrain dopamine neuron subtypes

To further explore the expression profiles of the DN-VTA and DN-SN populations in hMOs, we evaluated their similarity to established DN subtype markers from mouse, primate, and post-mortem human studies (Supplementary Table 1)^[3–9, 15, 20, 24, 31, 34, 37–39]^. We defined 3 DN marker groups; pan DN markers (expressed ubiquitously across midbrain DNs), preserved DN markers (higher expression in surviving DNs in PD), and vulnerable DN markers (higher expression in DNs lost in PD) (Fig.2C). Both DN-VTA and DN-SN clusters expressed pan DN markers including *TH* and DN transcription factors *FOXA1, FOXA2, LMX1A,* and *LMX1B* (Fig.2C). As expected, the DN-VTA cluster had higher expression of genes known to be upregulated in the VTA relative to the SN (bold genes in the preserved DN marker group of Fig.2C), including *ADCYAP1, NEUROD6,* and *CALB2.* We also observed high expression of vesicular glutamate transporter *SLC17A6* and a low expression of glutamate decarboxylase 1 and 2 (*GAD1* and *GAD2*) in the DN-VTA population, characteristic of previously defined glutamate and GABA co-releasing DNs found in the VTA ^[19, 34, 40, 41]^ (Fig.2C,D). In addition, DN-VTA cells have high expression of defined resilience genes expressed in surviving DNs in several PD mouse models and PD human postmortem tissue studies, consistent with their VTA-like identity, including *POU3F1*, *SORCS3*, and *TMEFF2*^[7, 9, 15, 20]^ (non-bold genes in the DN marker group of Fig.2C). Conversely, the DN-SN population has high *SOX6* expression classically used to distinguish DNs of the SNpc from those found in the VTA. Additionally, expression of SN markers, *ALDH1A1* and *NDNF,* and DN vulnerability marker, *ANXA1,* were upregulated in the DN-SN cluster^[3, 4, 8, 9, 15, 20, 34, 38]^ (Fig.2C,D). *ANXA1* is established as a marker of DNs found in the ventral tier of the SN co-expressing *SOX6* and *ALDH1A1*, and are shown to be the most vulnerable to degeneration in PD. We also observe an increase in genes associated with oxidative stress response in the DN-SN population consistent with increased basal oxidative stress in DNs of the SNpc relative to the VTA (Fig.2D). Conversely, an increase in genes associated with synaptic activity are observed in the DN-VTA population similar to previous reports^[20, 42]^(Fig.2D).

### Selective vulnerability of dopamine neurons occurs over time in *SNCA* triplication midbrain organoids

To validate the specific loss of the DN-SN but not the DN-VTA population observed in *SNCA* Trip hMOs, we performed immunofluorescent staining of pan DN marker TH, and broad DN-VTA and DN-SN subtype markers CALB1 and ALDH1A1 respectively (Fig.3A). We show that the total TH*+* area is significantly reduced in *SNCA* Trip relative to Iso CTL hMOs at 5 months, consistent with our scRNAseq data. The reduction is not observed at an earlier timepoint (3 months old) (Fig.3B). Quantification of the ratio of TH+ cells that also express CALB1 revealed no changes in the proportion of TH+CALB1+ neurons between *SNCA* Trip and Iso CTL hMOs at either 3 months or 5 months (Fig.3C). However, the proportion of TH+ cells co-expressing ALDH1A1 was significantly reduced in *SNCA* Trip hMOs relative to Iso CTL hMOs at 5 months old, but not earlier (Fig.3D). These results confirm that the lost TH*+* DNs are predominantly those expressing ALDH1A1, defining the DN-SN cluster in our scRNAseq dataset. Taken together, we show that our defined DN-VTA and DN-SN neuron populations in hMOs resemble SN and VTA DN populations described in the mouse, primate, and human brain datasets, with increased vulnerability of ALDH1A1*+* DN-SN neurons in *SNCA* Trip hMOs.

**Figure 3.**
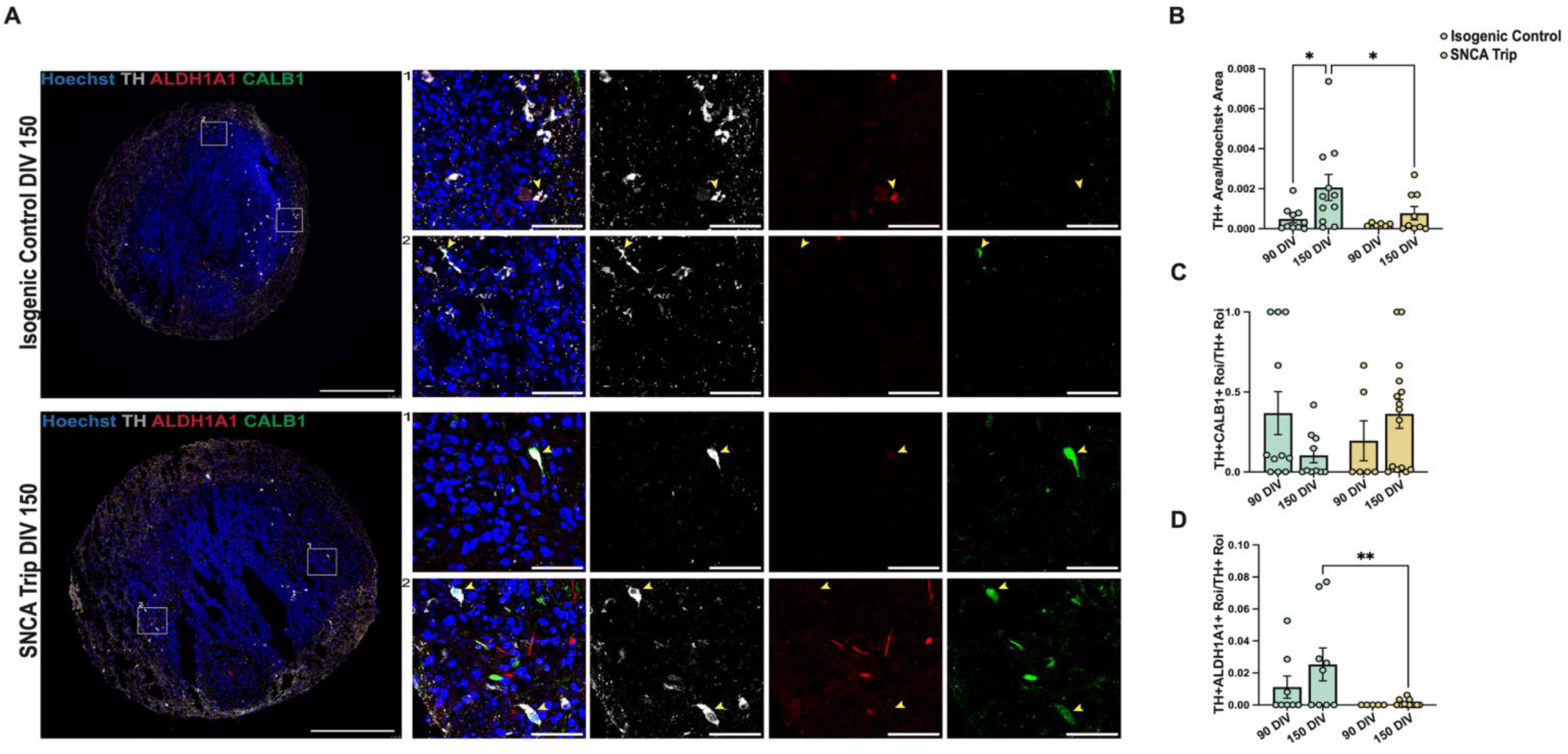
Immunofluorescent validation of the loss of ALDH1A1+ dopaminergic neurons in *SNCA* triplication midbrain organoids. **(A)** Representative image of an Iso CTL and *SNCA* Trip hMO at 150 days stained against TH (white), ALDH1A1 (red), and CALB1 (green). Representative close-up images highlight TH+ALDH1A1+ or TH+CALB1+ cells. Scale bar on whole organoid images is 500µm. Scale bar on close up representative images is 50µm. **(B)** Quantification of total TH+ area normalized to Hoechst area in Iso CTL and *SNCA* Trip hMOs at 90 days and 150 days. (n = 5-12 hMOs per condition, N=2). **(C)** Quantification of the proportion of TH+ cells that are also CALB1+ in Iso CTL and *SNCA* Trip hMOs at 90 days and 150 days. (n = 5-12 hMOs per condition, N=2). **(D)** Quantification of the proportion of TH+ cells that are also ALDH1A1+ in Iso CTL and *SNCA* Trip hMOs at 90 days and 150 days. (n = 5-12 hMOs per condition, N=2).

### Dopamine neuron population diversity and selective vulnerability is conserved across midbrain organoids from sporadic Parkinson’s disease patients

To investigate whether the DN-SN and DN-VTA populations, along with their differential vulnerability, are conserved across hMOs derived from different genetic backgrounds, we generated hMOs using independent iPSC lines from 4 healthy control (CTL) individuals and 3 sporadic PD (SP-PD) patients. To maximize our ability to capture disease relevant mechanisms prior to cell loss, we preformed scRNAseq on hMOs after 100 days and 200 days in culture. Using the same processing pipeline as with the *SNCA* Trip and Iso CTL hMO dataset, we merged our 100-day and 200-day datasets from all CTL and SP-PD hMOs and identified 14 distinct cell populations based on their expression of characteristic cell type markers and inferred transcription factor activity via SCENIC analysis (Fig.4A-B) (Supplementary Fig.2A). Notably, we were able to identify two DN populations resembling the DN-SN and DN-VTA populations identified in our *SNCA* Trip and Iso CTL hMOs. We observed expression of pan DN markers (*TH, NR4A2*) in both populations, *ALDH1A1* in the DN-SN population, and *SLC17A6* and *CALB2* in the DN-VTA population (Fig.4B). Both DN populations showed increased inferred activity of dopamine specific transcription factors similar to the *SNCA* Trip and Iso CTL dataset (Supplementary Fig.2B-F). To further characterize the DN-SN and DN-VTA clusters in the SP-PD and CTL dataset we looked at the expression of the defined DN subtype markers previously used (Fig.2C). Similar to our *SNCA* Trip and Iso CTL hMO dataset, both DN-SN and DN-VTA clusters express pan DN markers, with increased expression of preserved DN markers including *CALB1*, *CALB2*, and *SLC17A6* in the DN-VTA cluster, and vulnerable DN markers including *SOX6*, *ALDH1A1*, and *ANXA1* in the DN-SN cluster (Fig.4C).

**Figure 4.**
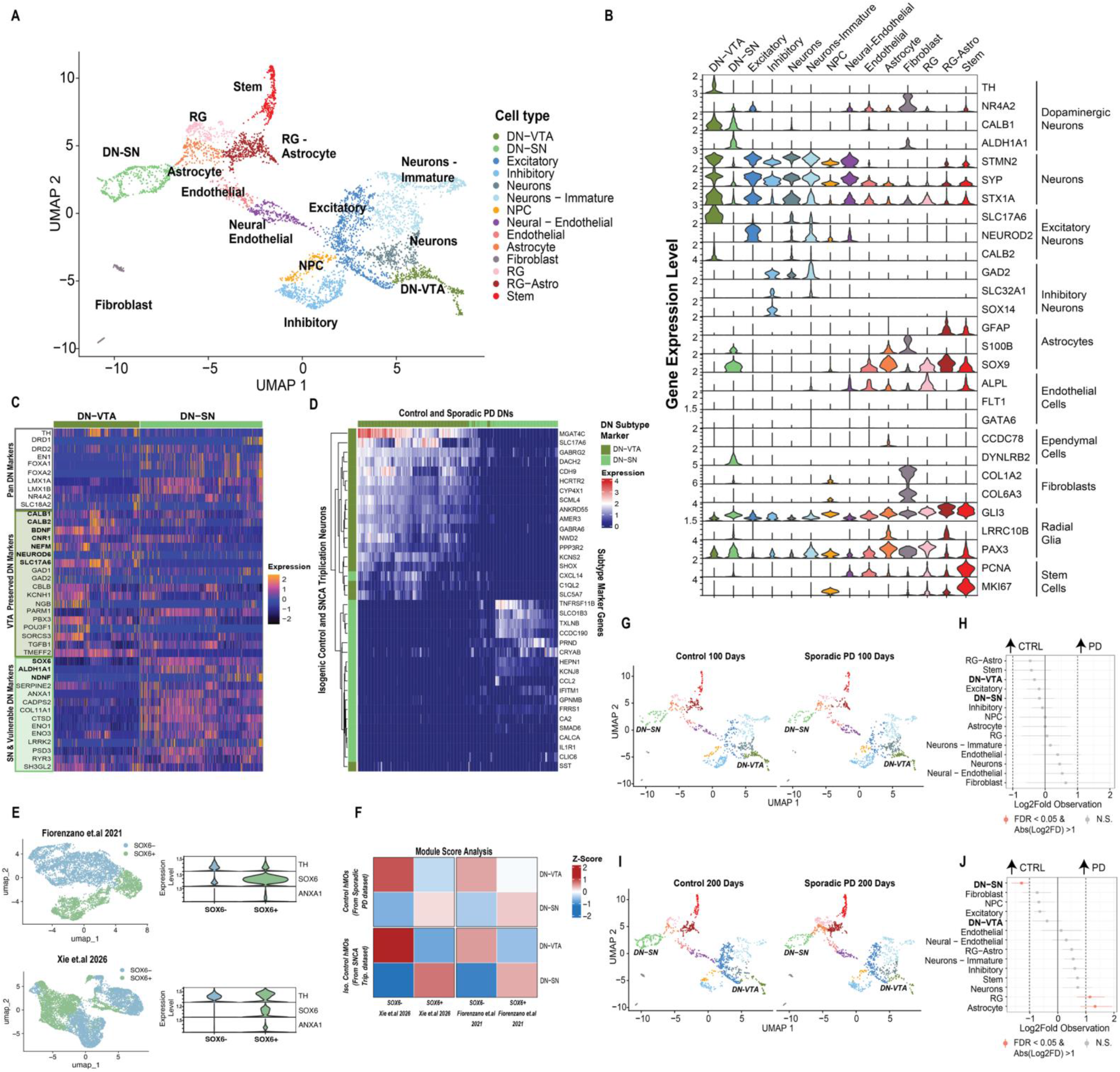
Single cell transcriptomic characterization of cell types in sporadic Parkinson’s disease and control midbrain organoids. **(A)** UMAP plot showing annotated clusters derived from all sporadic PD and control hMOs at 100 days and 200 days. **(B)** Violin plot of canonical cell type markers for each cell cluster. DN-VTA and DN-SN are distinguished by ALDH1A1 expression in DN-SN cells. **(C)** Heatmap of genes defined in previous studies as either pan dopaminergic (expressed across VTA and SN dopaminergic neurons), highly expressed in VTA and preserved dopaminergic neurons in PD, or highly expressed in SN and susceptible dopaminergic neurons in PD. Expression of selected genes are shown for DN-SN and DN-VTA clusters. **(D)** Hierarchical clustering of DN-SN and DN-VTA cells from *SNCA* Trip and Iso CTL hMOs based on the top 20 differentially expressed genes from DN-SN and DN-VTA cells derived from SP-PD and CTL hMOs at 200 days. **(E)** Left: UMAP plots of dopaminergic neurons generated using published data from midbrain organoids, clustered and annotated based on SOX6 expression. Right: Violin plot validating SOX6 and ANXA1 expression in SOX6- and SOX6+ clusters. **(F)** Average expression of top 100 differentially expressed genes from SOX6- and SOX6+ clusters used in (E) compared to DN-SN and DN-VTA clusters described in both our datasets. **(G)** Split UMAP of SP-PD and CTL hMOs at 100 days. **(H)** Permutation test comparing cell type proportions between SP-PD and CTL hMOs at 100 days. Significant changes in cell proportion are described as those with FDR < 0.05 and log2 fold change >1. **(I)** Split UMAP of SP-PD and CTL hMOs at 200 days. **(J)** Permutation test comparing cell type proportions between SP-PD and CTL hMOs at 200 days.

We next compared the transcription levels of genes defining the DN-SN and DN-VTA clusters in the SP-PD and CTL dataset to those in the *SNCA* Trip and Iso CTL datasets. For this, we performed differential gene expression (DGE) analysis between DN-SN and DN-VTA cells in the SP-PD and CTL dataset, took the top 20 differentially expressed genes (DEGs) in both clusters, and used these genes to perform unbiased hierarchical clustering of DN-SN and DN-VTA cells in the *SNCA* Trip and Iso CTL dataset. We found that the DN-SN and DN-VTA cells in the *SNCA* Trip and Iso CTL dataset generally cluster into the expected DN subtypes based on expression of the cluster markers taken from the SP-PD and CTL dataset (Fig.4D). This points to transcriptional similarity in key subgroup defining genes between the two datasets, increasing confidence in our DN subtype annotations between datasets.

To further compare the DNs observed in our datasets with those described in published hMO data, we leveraged publicly accessible scRNAseq datasets from two separate hMO studies, isolated the healthy control DNs, and re-clustered and annotated them based on *SOX6* expression (Fig.4E)^[24, 42]^. We then compared the transcription levels of the top 100 expressed genes in *SOX6-* and *SOX6+* populations from both public datasets independently to the DN clusters (DN-VTA and DN-SN) from each of our datasets. The average expression of each list of 100 genes was computed per cell in both of our DN clusters and compared to the expression level of a set of random control genes. We observed high gene expression in our DN-SN and DN-VTA populations to the top 100 expressed genes from the *SOX6+* and *SOX6-*populations, respectively, from both public datasets (Fig.4F). This analysis was done on isolated control neurons from our *SNCA* Trip and SP-PD datasets to remove potential disease associated signatures in the PD lines that may be absent in the reference control datasets. These results indicate that the DN-SN and DN-VTA populations endogenously develop in hMOs derived across distinct genetic backgrounds, with conserved transcriptional similarity of the distinct DN populations across hMOs derived using both our in-house protocol and separate protocols in other labs^[24, 42, 43].^

Finally, to determine if there were any changes in cell type proportions between the CTL- and SP-PD-hMOs over development, we performed permutation tests between both groups at each time point. We see no significant changes in the proportion of cell clusters at 100 days (Fig.4G-H), however we observe a significant reduction in DN-SN cells in SP-PD hMOs at 200 days (FDR = 0.002, Log2FD = -1.28) (Fig.4I-J), similar to our previous observations in *SNCA* Trip hMOs relative to their Iso CTLs (Fig.1E). We also observed a significant increase in the RG and Astrocyte clusters in SP-PD hMOs relative to CTLs (Fig.4J). Taken together, these results suggest that the clusters identified as DN-VTA and DN-SN in the SP-PD and CTL hMOs resemble those identified in the *SNCA* Trip and Iso CTL hMOs, with conserved vulnerability of the DN-SN cluster in hMOs derived from different PD patients.

### Vulnerable dopamine neurons in midbrain organoids derived from Parkinson’s disease patients display increased expression of mitochondria-related transcripts

We next characterized transcriptional changes between DN-SN neurons derived from PD hMOs and CTL hMOs to identify genes that may contribute to increased vulnerability in the DN-SN population. Since all DN-SNs were lost in the *SNCA* Trip hMOs, we could not make this comparison using the *SNCA* Trip background. However, as a proportion of DN-SNs survived in the SP-PD hMOs at 200 days, we were able to perform DGE analysis between SP-PD and CTL DN-SNs at this timepoint, and at 100 days, prior to DN loss. DGE analysis at 100 days revealed 323 significantly upregulated and 161 significantly downregulated (average Log2FC > 0.5 or <-0.5, adjusted pVal <0.001) genes in the SN-DNs derived from SP-PD hMOs relative to CTL hMOs (Supplementary Fig.3A). Gene ontology (GO) over representation analysis (ORA) (using the biological processes and molecular function databases) of the genes downregulated in DN-SN cells of SP-PD hMOs revealed terms involved in metabolic processes and microtubule formation. However, no significant pathway enrichment was observed in the upregulated genes. (Supplementary Fig.3A). In contrast, DGE analysis at 200 days revealed a larger number of DEGs bidirectionally, with 3495 significantly upregulated and 1539 significantly downregulated genes in DN-SNs derived from SP-PD relative to CTL hMOs (average Log2FC > 0.5 or <-0.5, adjusted pVal <0.001) (Fig.5A). Interestingly, ORA revealed multiple upregulated terms involving mitochondrial gene expression, mitochondrial complex chain formation and activity, and protein translation, with cell-cell adhesion being the only downregulated term (Fig.5B). These results suggest potential mitochondrial dysfunction, increased mitochondrial biogenesis, and/or increased translation of mitochondrial genes in surviving DN-SN cells derived from SP-PD hMOs.

**Figure 5.**
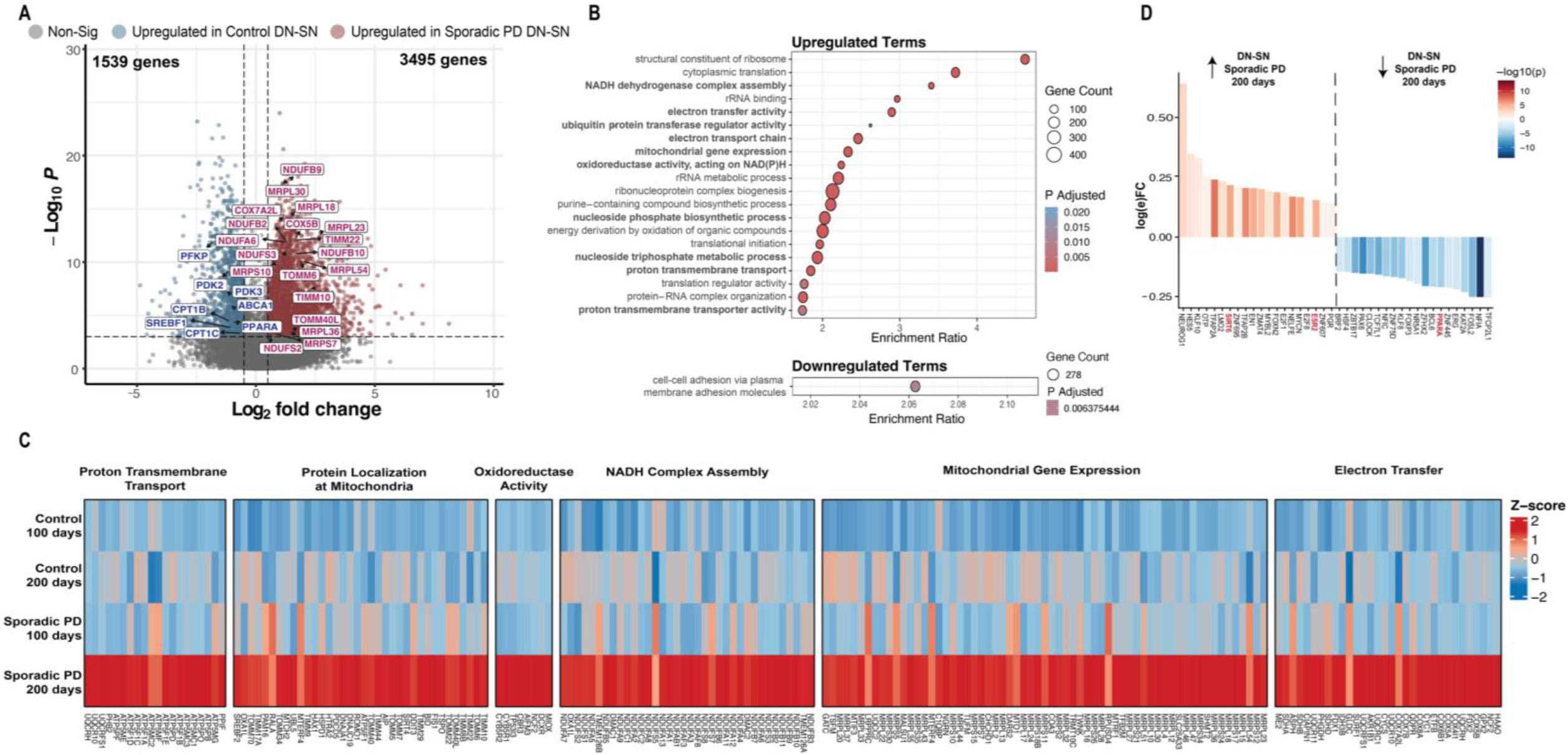
Differential gene expression analysis between Parkinson’s disease and control midbrain organoids in DN-SN cells indicates changes in mitochondrial gene expression. **(A)** Volcano plot highlighting differentially expressed genes between SP-PD and CTL DN-SN cells at 200 days. Selected mitochondria related genes are labelled (FDR < 0.001 and log2 fold change > 0.5 or < -0.5). **(B)** Gene ontology over representation analysis of significantly up- and down-regulated genes between SP-PD and CTL DN-DN cells reveals multiple mitochondria related pathways upregulated in SP-PD hMOs at 200 days. **(C)** Heatmap showing relative expression of mitochondrial genes from the mitochondria related pathways described in (B) between SP-PD and CTL DN-SN cells from both 100 days and 200 days. **(D)** SCENIC analysis showing significantly up- and down-regulated regulons between SP-PD and CTL DN-SN cells at 200 days.

We then compared the expression of genes enriched in mitochondrial GO terms across hMO development to determine if the observed increase in mitochondrial gene expression in SP-PD hMOs was unique to the 200-day timepoint. We observe an increase in mitochondria related genes in the DN-SN population from 200-day SP-PD hMOs relative to 100-day SP-PD hMOs, correlating increased mitochondrial gene expression with the timepoint in which DN-SNs are lost (Fig.5C, Supplementary Fig.3B). Similar analysis done on CTL hMOs revealed a less pronounced increase in mitochondrial gene expression at 200 days vs 100 days, with a single mitochondria related term in the upregulated genes (“mitochondrial gene expression”) (Supplementary Fig.3C). Investigation into the genes related to the “mitochondrial gene expression” term revealed an abundance of mitochondrial ribosomal genes, but not genes associated with mitochondrial complex chain formation and activity as seen in the SP-PD hMOs (Supplementary Data Table 10). These results suggest that changes in mitochondria related gene expression occur over time in healthy DN-SN cells, consistent with increased maturity and potential increases in bioenergetic demands. However, a pronounced increase in mitochondria genes relating to mitochondrial activity and respiratory chain complex assembly is observed in DN-SN cells from SP-PD hMOs between 100 days and 200 days of development. This increase is not present in CTL hMOs and correlates to the time in which DN-SNs are significantly lost in SP-PD hMOs. We also observed no changes in mitochondrial gene expression in the DN-VTA cluster between SP-PD and CTL hMOs at either timepoint (Supplementary Data Table 11-14). This suggests that DN-SN cells are specifically sensitive to changes in mitochondrial function relative to other cell types, with no changes in mitochondrial gene expression in preserved DNs.

We next preformed gene regulatory analysis using SCENIC on the DN-SN population to determine if we could identify regulatory networks associated with mitochondrial gene expression. SCENIC analysis, between SP-PD and CTL DN-SN cells at 200 days revealed an increase in activity of two transcription factors previously associated with positive regulation of mitochondrial genes, estrogen receptor 2 (*ESR2*) and the deacetylase sirtuin 6 (*SIRT6*) ^[44, 45]^ (Fig.5D). Both *ESR2* and *SIRT6* activity is increased in 200-day SP-PD compared to 100-day SP-PD hMOs, supporting an age dependent increase in mitochondrial gene expression (Supplementary Fig.3D). Interestingly, we observe a decrease in peroxisome proliferator-activated receptor alpha (PPARA) activity and gene expression (Fig.5A,D). PPARA is a co-activator of the transcription factor peroxisome proliferator-activated receptor gamma coactivator 1-alpha (PGC1*α*), and is involved in regulation of mitochondrial fatty acid oxidation gene expression^[46–48]^. As expected, we also see a downregulation in gene expression of multiple genes involved in fatty acid oxidation in the DN-SN population from SP-PD hMOs including *PFKP, PDK2, PDK3, ABCA1*, *CPT1B, CPT1C,* and *SREBF1*, consistent with reduced PPARA activity (Fig.5A). A reduction in mitochondrial fatty acid oxidation genes leads to reduced ATP production via impaired oxidative posphorylation^[49]^. Additionally, fatty acid oxidation has been linked to increased NADPH levels, resulting in reduced intracellular ROS and oxidative stress^[49]^. Taken together, these data nominate mitochondrial dysfunction via reduced fatty axid oxidation genes as a mechanism underlying cell vulnerability in DN-SN cells of SP-PD hMOs.

## Discussion

Although PD is defined by the progressive degeneration of DNs in the midbrain, not all DNs within this region are affected equally. Advances in, and increased availability of, scRNAseq technologies have enabled researchers to identify subpopulations of DNs within the broadly defined SNpc that contribute to distinct functional outputs and display differential susceptibility to PD^[15, 16, 34, 50]^. The mechanism underlying differential DN vulnerability in PD is not fully understood, but VTA DNs are hypothesized to be protected by their smaller axonal arbors, lower metabolic demands, and reduced reliance on Cav1.3 Ca²⁺ channels, hypothesized to increase intracellular Ca²⁺ and metabolic demand to buffer Ca²⁺ out of the cytoplasm^[51–53]^.

Furthermore, VTA DNs exhibit elevated expression of protective genes, including *BDNF* and the calcium-buffering gene *CALB1*, linking calcium buffering capacity to neuronal survival^[38]^. More recently, researchers have begun to dissect distinct subpopulations within the SNpc itself, revealing that ventral tier SNpc DNs are particularly susceptible to degeneration and are among the first to be lost as PD progresses^[15]^. Dorsal tier SNpc DNs, by contrast, are more resilient and more closely resemble VTA DNs, exhibiting higher *CALB1* expression^[3, 4, 15]^. Functional studies have linked ventral tier SNpc DNs to the control of motor acceleration, while dorsal tier DNs, like VTA DNs, are linked to motor deceleration and reward response^[8]^. These findings offer insight into the progression of motor symptoms. Bradykinesia, characterized by slowed movement, is among the earliest motor symptoms of PD, consistent with the early loss of the ventral tier DNs that support motor acceleration.

Numerous studies have since sought to characterize this vulnerable ventral DN population further. Markers such as *ALDH1A1* and *ANXA1* have been identified as differentially expressed in ventral tier DNs, establishing them as characteristic markers of vulnerable neurons^[3, 8]^. scRNAseq have enabled researchers to examine changes in gene expression within specific molecular pathways, nominating mechanistic drivers of DN vulnerability. These analyses have identified gene ontology pathways related to lysosomal function, oxidative stress, and mitochondrial activity as consistently upregulated in vulnerable DNs, while pathways associated with synaptic activity are generally downregulated relative to preserved DN populations^[15, 16]^. However, further work is needed to define the full extent of DN subtype diversity in the human brain, their differential susceptibility to PD, and the precise molecular contributors to this vulnerability. Addressing these questions will require human model systems that capture cellular diversity, including distinct DN subpopulations, in a disease-relevant context.

Brain organoids are an emerging model commonly used for the study of neurological diseases, with recent studies showing an increasing ability of brain organoids to capture complex disease relevant phenotypes^[26, 28, 29, 54, 55]^. Remarkably, these studies highlight the capacity of iPSC derived brain organoids to model various aspects of neurodegenerative disease pathology. For example, PD brain organoid models show an accumulation of phosphorylated α-synuclein, loss of TH+ cells, and electrophysiological aberrations^[26, 28, 29, 54, 55]^. Previous studies have suggested the existence of multiple DN subpopulations in hMOs, resembling preserved and vulnerable DN populations, based on transcriptional identity and protein expression^[24, 42, 56]^. However, these studies failed to demonstrate the selective loss of “vulnerable” DNs in PD hMO models^[3, 15, 20]^. Establishing the capacity of hMOs to capture selective vulnerability of DNs in PD would be an important step to validate their use in studying nuanced PD-associated mechanisms related to DN loss.

Here we describe the presence of two DN subpopulations resembling preserved DNs of the VTA and vulnerable DNs of the SN in hMOs across multiple cell lines. Importantly, we show, for the first time, that DNs resembling those found in the SN are selectively lost in PD patient derived hMOs with no loss of DNs resembling those found in the VTA. Remarkably, the loss of DN-SNs was conserved across hMOs derived from multiple PD patient lines, including both familial and sporadic PD cases. DGE analysis between remaining DN-SN cells from PD vs CTL hMOs revealed a significant increase in mitochondrial electron transport chain related gene expression, but a decrease in mitochondrial fatty acid oxidation genes. The reduction in fatty acid oxidation genes correlates with the reduction in *PPARA* gene expression and inferred regulon activity, a known regulator of fatty acid oxidation gene transcription. In contrast, it has been shown that the activation of PGC1*α* through other co-activators such as NRF-1, NRF-2, ERR*α*, SIRT6, and ESR2, controls transcription of genes related to electron transport chain activity and oxidative phosphorylation ^[47]^. Thus, the observed increase in expression of mitochondrial electron transport chain and oxidative posphorylation genes may be linked to ESR2 and SIRT6 activity, potentially as a compensatory mechanism for decreased expression of *PPARA* and fatty acid oxidation genes. These results link mitochondrial dysfunction to DN-SN cell vulnerability observed in PD hMOs. Here, mitochondria impairment may be driven by reduced PPARA activity and fatty acid oxidation gene expression which regulates efficient ATP production in mitochondria. Briefly, fatty acid chains are converted into acyl-CoA which gets shuttled into the mitochondrial matrix and enters the TCA cycle, producing nicotinamide adenine dinucleotide (NADH), driving ATP generation through oxidative phosphorylation^[49]^. This process if regulated through fatty acid oxidation genes controlled by PPARA, therefore, reduced fatty acid oxidation gene expression may lead to impaired break down of fatty acid chains into acyl-CoA, reduced acyl-CoA shuttling into the mitochondria, and reduced ATP production, a process which could be detrimental to DNs which have high energy demands. Though compelling, it would be necessary to validate this hypothesis through quantification of ATP production and manipulation of the fatty acid oxidation pathway directly in hMO DNs.

Changes in mitochondrial gene expression have consistently been observed in PD models^[15, 31, 57–59]^. It has been well documented that chemicals interfering with mitochondrial respiratory chain activity lead to parkinsonism^[60–66]^. Furthermore, postmortem PD tissue analyses have reported changes in Complex I, Complex II, and Complex IV activity and expression^[67–70]^. These reports highlight the importance of mitochondrial function in DN survival and PD progression, further supported by decreased mitochondrial activity being commonly observed in sporadic PD patients, without known mutations in mitochondria related genes^[67, 68, 71]^. However, the directionality of mitochondrial gene expression varies between studies. Some show a downregulation of mitochondrial genes in vulnerable PD DNs ^[57, 72, 73]^, whereas others show an upregulation of related genes^[15, 31, 42]^ alongside a simultaneous decrease in PPARA activity similar to our findings^[15]^. The lack of consensus between studies may be due to inconsistencies in the disease timepoints analyzed. Therefore, it is possible that vulnerable DNs display an early increase in mitochondrial gene expression as a compensatory stress response to elevated bioenergetic demands prior to large-scale pathology, followed by reduced gene expression at DN end-stages. It is important to note that our observed increase in mitochondria gene expression is biased towards surviving DNs. Based on these experiments, we cannot conclude if the same increase in mitochondria gene expression would be observed in lost DNs prior to their loss. To answer this, it would be important to characterize DN-SNs across multiple timepoints in PD hMO development to track changes in mitochondrial gene expression during DN-SN degeneration. Furthermore, since we could not analyze the DN-SN population in *SNCA* Trip hMOs, it will be essential to assess mitochondrial gene expression, and mitochondrial function, across disease progression in multiple cell lines to determine whether this represents a conserved pathway of neuronal vulnerability or if distinct PD cases drive divergent mechanisms of vulnerability.

Regardless, it is remarkable that selective vulnerability of DNs is captured in hMO models of PD, suggesting conserved developmental and susceptibility differences between DN subpopulations in *in vitro* hMO and *in vivo* models. Important future directions stemming from this work include enriching DNs from hMOs to gain a deeper understanding of the subtypes of *SOX6*+ and *CALB1*+ DNs that may be present in hMOs. Recent work has identified 20 subpopulations of DNs in the mouse midbrain with distinct projection targets and functional roles^[50, 74]^ adding to the complexity of the DN system. In future experiments, it would be important to verify and investigate DN subpopulations with increased granularity in hMOs, which may include the use of more complex systems such as midbrain-striatal assembloids to assess diversity in projection patterns and functional activity. Progress in our understanding of the mechanisms underlying selective vulnerability in PD has been limited by the lack of translatability of animal models and reliance on end-stage PD postmortem tissue where vulnerable DNs are already lost. Thus, these experiments highlight the significance of hMOs to capture disease pathology including the selective vulnerability of DN-SN cells while revealing diseases relavant mechanisms associated with their vulnerability. Therefore, as shown here, hMOs are of particular interest as these results present a novel opportunity to study PD relevant changes in vulnerable DN populations across disease progression in relevant cell types derived from patients, opening the door to patient stratification and future personalized therapies.

## Methods

### iPSC maintenance and hMO generation

The use of human iPSCs and iPSC-derived cells in this research was approved by the McGill University Research Ethics Board. All study procedures adhered to the Declaration of Helsinki. SNCA Trip and Iso CTL iPSCs were generated and provided by Dr. Tilo Kunaths laboratory in the University of Edinburgh^[75, 76]^ . All iPSCs (Supplementary Table 2) were thawed from liquid nitrogen in a 37°C water bath, plated on a Matrigel coated dish in mTeSR media, and supplemented with 10µM Y-27632. mTeSR was changed daily without Y-27632 and iPSCs were checked for contamination and differentiation. Differentiated cells were scraped off prior to media change, 70-80% purity was maintained at all times. iPSCs were passaged once they reached 60-70% confluency, and a minimum of two passages were done prior to differentiation. *SNCA* Trip and Iso CTL hMOs were generated using the protocol described in Mohamed and colleagues^[77]^, while SP-PD and CTL hMOs were generated using the more recent, modified, protocol described in Mohamed and colleagues^[43]^.

### hMO dissociation and scRNAseq preparation

All hMOs were dissociated with a combination of enzymatic digestion and mechanical dissociation to prepare single cell suspensions, FACS sorting was used to enrich for live cells and samples were processed for scRNAseq using either 10X Genomics droplet sequencing or ParseBio split-seq technology followed by NextGen Illumina sequencing. *SNCA* Trip and Iso CTL individual hMOs were pooled together (3-4 hMOs) for each sample in one 15mL tube and washed three times with Dulbecco’s PBS (D-PBS) to completely remove remaining culture media. hMOs were incubated in 2mL of TrypLE express without phenol red at 37°C for a total of thirty minutes. Mechanical dissociation by pipette trituration (slowly pipetting up and down ten times) was performed three times after ten minutes of incubation. Afterwards, 8mL of D-PBS was added to the samples to stop the enzymatic reaction. The samples were filtered through a 30µm filter (Miltenyi Biotec) to remove any clumps remaining after digestion and dissociation.

The cells were centrifuged for 5 minutes and washed twice in D-PBS + 0.1% BSA. Single cell suspensions were incubated with (live/dead stain) then sorted for live cells using FACSAria Fusion (Becton-Dickinson Biosciences). After manual counting, the single cell suspensions were diluted to 1000 cells/ml targeting and loaded into Chromium NextGEM Chip G targeting ∼5,000 cells for sequencing using the Chromium Next GEM Single Cell 3’ reagent kit v3.1 and run on the 10X Chromium Controller for GEM creation. All proceeding thermocycler steps were carried out on a Bio-Rad C1000 Touch thermal cycler. Following GEM-RT incubations, samples were stored at 4°C overnight. Post GEM-RT cleanup and cDNA amplification were carried out per manufacturer protocol. Samples were stored at -20°C until they were processed for library generation. 3’ gene expression and cell surface protein libraries were constructed per manufacturer protocol and stored at -20°C until sequencing submission. 25 mL of each sample library was sent for next generation sequencing at the McGill Genome Centre and using the NovaSeq6000 sequencer. For the sporadic PD and control lines 10-20 hMOs were pooled together into one 50mL tube, then washed three times with Dulbecco’s PBS (D-PBS) (Wisent) to completely remove remaining culture media before transfer to a gentleMACS M-Tube. The D-PBS was completely removed and 2mL of TrypLE express without phenol red was added to each sample. The hMOs inside the M-Tube were next placed on an automated the GentleMACS Octo Heated dissociation device. The settings for the dissociation were as follows: 37°C is ON. Spin 20rpm for 24 minutes. Spin 197rpm for 1 minute. After GentleMACS dissociation, 8mL D-PBS was added to the samples to stop the enzymatic reaction. The samples were filtered through a 30µm filter (Miltenyi Biotec) to remove any clumps remaining after digestion and dissociation. The samples were then washed twice more with D-PBS.

### scRNAseq data processing and quality control

Raw sequencing files were processed using the automated scRNAbox pipeline^[78]^. FASTQ files from 10X genomics were reprocessed using an updated version of the CellRanger pipeline (v5.0) with intron inclusion selected and optimized STAR alignment configuration. ParseBio FASTQ sequencing libraries were processed with the Split-pipe pipeline installed on Digital Computing Alliance following the ParseBio directions for demultiplexing samples to generate aligned genes, barcodes, and expression matrixes.

The features, barcodes, and raw counts expression matrices were combine using Seurat v5 in R. For all samples the following quality control filters were applied to maintain high confidence single cells; total RNA counts < 20, unique RNA counts > 500 and < 10,000, and percent mitochondrial reads < 20%. Remaining potential doublets were removed using doublet finder. The feature matrixes were then filtered to remove non-coding RNA and mitochondrial DNA transcripts then denoised and imputed using the ALRA algorithm implemented in R^[30]^ and used for downstream analysis unless otherwise indicated.

### scRNAseq Cell Clustering

Clustering and annotation analysis was performed using Seurat v5 functions using R scripts. For each dataset biological samples were integrated using the Seurat’s classic Canonical Correlation Analysis workflow. The *FindVariableFeatures* function was used to select the top 5000 most variable genes for cluster analysis from the normalized ALRA counts matrix. The 5000 genes were normalized and scaled prior to linear dimensional reduction by principal component analysis (PCA). Elbow plots and JackStraw tests were used to determine the number of principal components (PCs) to keep. A range of k values for the Shared Nearest Neighbor (SNN) graph and resolutions for leiden network detection were tested using the *FindClusters* function. Cluster branching across resolutions was analyzed using the clustree R package^[79]^ and visually inspected by Uniform Manifold Approximation and Projection (UMAP). Optimal parameters were selected to generate clusters for annotation. For the SNCA trip and CTL dataset the following parameters applied were: 25 PCs, 25 nearest neighbours (k.param) and resolution of 0.75. For the sporadic and health lines dataset the parameters were: 20 PCs, 35 k.param and resolution of 0.5.

To annotate clusters by cell type three approaches were used. 1) Differential gene expression analysis using *FindAllMarkers* to find upregulated genes for each cluster. Genes with adjusted p-values < 0.01 and Log2FoldChange > 0.25 were considered significant and selected as cluster markers. The package EnricheR was then used to match the cluster markers to cell type libraries (Azmuth*CellTypes 2021, Allen Brain* Atlas *10x* scRNA *2021, CellMarker*) and Gene Ontology pathways (*Biological* Process*2023, CellularComponent, Molecular Function, 2023*). 2) Cell type predictions were mapped using the Seurat data mapping annotation workflow from multiple difference single cell reference datasets listed in (Supplementary Table 3). 3) Module scores were calculated from a curated cell type marker list (Supplementary Table 4). The consensus of all three cell type predictions was used to determine the final cell type annotations.

### Gene regulatory network analysis using SCENIC

After scRNAseq data processing, quality control, and cell type annotation, we used the SCENIC pipeline (https://scenic.aertslab.org/) to reinforce annotations and infer disease-associated active gene regulatory network (GRN) elements. We saved expression matrices and metadata from our Seurat object and converted them to loom files with Loompy (https://linnarssonlab.org/loompy/). We then ran SCENIC analysis on Digital Research Alliance of Canada’s Rorqual cluster using the pySCENIC implementation (https://pyscenic.readthedocs.io/en/latest/installation.html#command-line-interface) through a docker image (aertslab/pyscenic:0.12.1) and with default settings. In short, GRN inference is done using grnboost2, pySCENIC predicts regulons, and cells’ regulon activity is scored with AUCell. The resulting regulons and their cell-level scores were added back to the Seurat object as metadata and downstream analysis proceeded in R.

### Cryosectioning and immunofluorescence staining of hMOs

hMOs were fixed in 4% PFA in PBS at 4°C overnight then switched to a 20% sucrose in PBS solution for 3 days to dehydrate tissue. hMOs were then placed in cryomolds, all remaining liquid was aspirated, and hMOs were embedded in OCT compound (Fisher HealthCare) and frozen over liquid nitrogen. Cryomolds were kept at -80°C until sectioning. Sectioning was done using a cryostat (ThermoFisher, CryoStar NX70 Cryostat) at a thickness of 10µM. For staining, slides were rehydrated in PBS for 15 minutes, the Super Pap Pen Liquid Blocker (IHCWorld, SPM0928) was used to draw a barrier around sections to prevent buffers from leaking off the slide. Following rehydration, sections were blocked (5% NGS, 0.05% BSA, 0.2% Triton-X in PBS) for 1 hour at room temperature followed by primary antibodies (Supplementary Table 5) diluted in blocking buffer overnight at 4°C. Sections were then washed in PBS 3 x 5 minutes and incubated in secondary antibodies diluted in blocking buffer for 1 hour at room temperature. Sections were washed again in PBS 3 x 5 mins and incubated in Hoechst (1:5000 diluted in PBS) for 10 minutes at room temperature, followed by a final set of washes in PBS 2 x 5 mins. Sections were then sealed with glass coverslips using AquaMount solution and imaged at 40X using the Leica TCS SP8 Confocal microscope. All images were analyzed using CellProfiler for detection of staining ROIs, and measurements of area and colocalization.

## Statistical analysis

All statistical analysis for immunofluorescent experiments was performed using GraphPad Prism version 10.4.1 for MacOS, GraphPad Software, Boston, Massachusetts USA, www.graphpad.com. For analysis of total TH+ cells and subtype proportions (Fig.2B-D), a Two-Way ANOVA with multiple comparisons was used. DGE analysis was done using Seurats V5.4 FindMarkers function with MAST testing. Significant differentially expressed genes were defined as those with Padj < 0.05 and Log2 Fold Change >0.5 for upregulated genes or <-0.5 for downregulated genes. Gene ontology analysis was done using WebGestalts online toolkit (https://www.webgestalt.org) with a Weighted Set Cover analysis (FDR <0.05). For permutation tests, the scProportionTest library was used in R Studio with an FDR cutoff of <0.05. Log2FD thresholds were set as 2 standard deviations away from the mean of the observed Log2FD values for all cell types within a given dataset. Calculated Log2FD thresholds were then rounded to the nearest whole value.

## Data and Code Availability

The processed and annotated scRNAseq data objects used in this manuscript is deposited in the UCSC Cell Browser: https://midbrain-organoid-aldh1a1.cells.ucsc.edu

Source Data are provided with this paper as “Supplementary Data Tables”.

## Acknowledgments

We would like to thank Carol X.-Q. Chen for helping with iPSC quality control and facilitating access to the iPSC lines. We also want to thank The Neuro Microscopy Core Facility for confocal microscope management, maintenance, and training, and The Neuro FACS platform for help in tissue dissociation and processing. We would like to thank the Hospital for Sick Children and Axol Bioscience for their help in reprogramming and karyotyping iPSC lines. This work was supported by a *Fonds d’Accéleration des Collaborations en Santé* grant from CQDM/MEI (Fon-FACS-013) and a Michael J. Fox Foundation grant (MJFF-021629). E.A.F. is supported by a Canada Research Chair (Tier 1) in Parkinson’s disease. The funders were not involved in study design, data collection and analysis, or the writing of this manuscript.

## Author Contributions

The project was conceived by G.D., R.A.T, and E.A.F. G.D. generated, cultured, processed, imaged and analyzed hMOs used for immunofluorescent analysis, analyzed scRNAseq data, and generated manuscript figures. R.A.T. processed raw scRNAseq data and prepared data objects, ran statistical analysis, helped in the generation of manuscript figures, and provided conceptual support for the manuscript. J.G. generated SCENIC scripts and aided in SCENIC analysis. T.M.G. processed hMO tissue for scRNAseq. P.L. generated and maintained hMOs used for scRNAseq experiments. J.S. aided in hMO tissue processing and dissociation for scRNAseq. M.M. aided in the generation and maintenance of hMOs. V.O. aided in hMO tissue preparation and processing for scRNAseq. Data were interpreted by G.D., R.A.T. and E.A.F. The manuscript was written by G.D. and E.A.F. with contributions by R.A.T., and T.M.D. The project was supervised by E.A.F.

