## Supplementary Figures and Tables for "Transcriptomic profiling of Parkinson’s disease patient-derived midbrain organoids reveals selective vulnerability of ALDH1A1+ dopamine neurons"

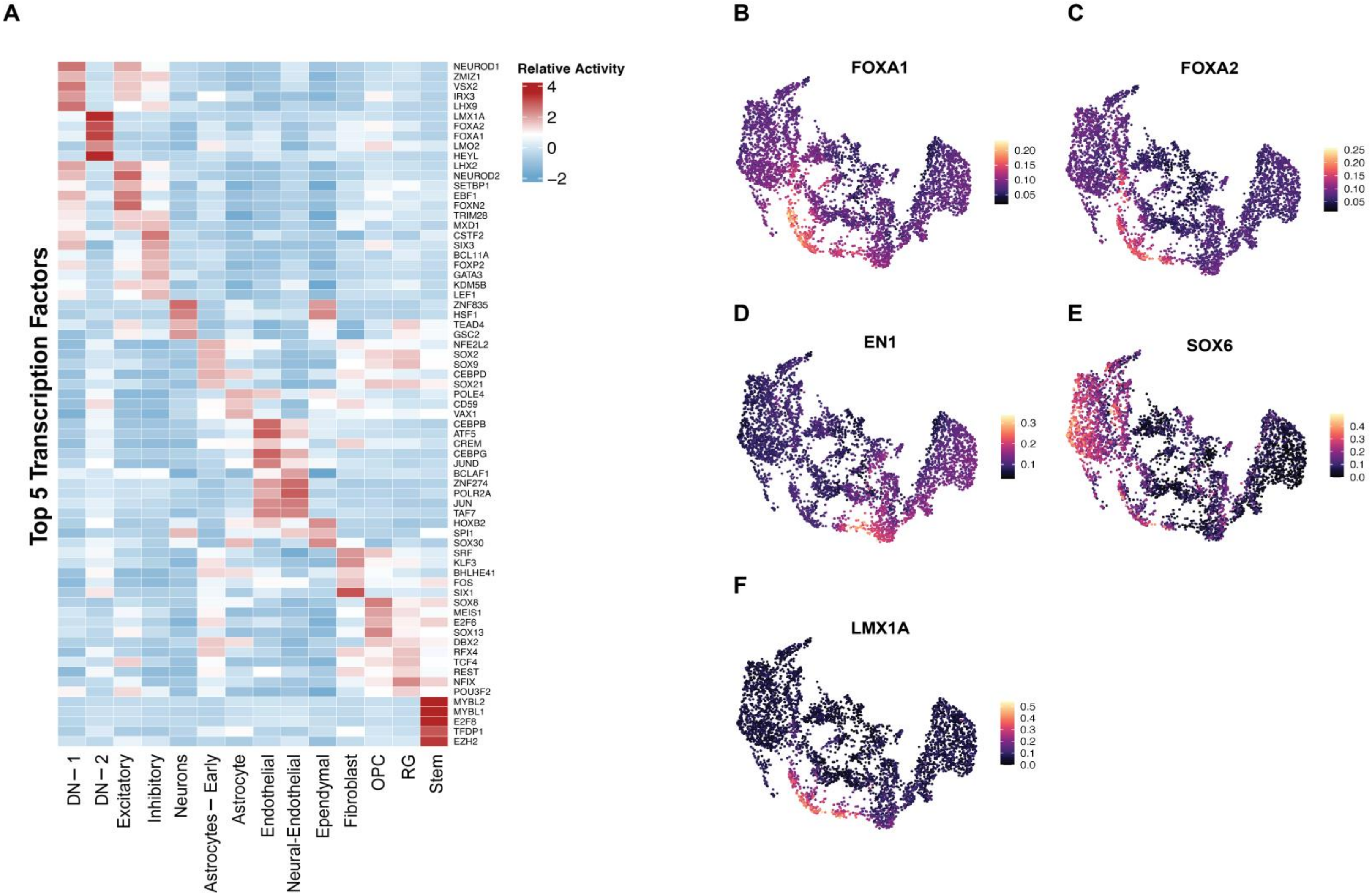

A
Top 5 Transcription Factors
Relative Activity
4
2
0
−2
NEUROD1
ZMIZ1
VSX2
IRX3
LHX9
LMX1A
FOXA2
FOXA1
LMO2
HEYL
LHX2
NEUROD2
SETBP1
EBF1
FOXN2
TRIM28
MXD1
CSTF2
SIX3
BCL11A
FOXP2
GATA3
KDM5B
LEF1
ZNF835
HSF1
TEAD4
GSC2
NFE2L2
SOX2
SOX9
CEBPD
SOX21
POLE4
CD59
VAX1
CEBPB
ATF5
CREM
CEBPG
JUND
BCLAF1
ZNF274
POLR2A
JUN
TAF7
HOXB2
SPI1
SOX30
SRF
KLF3
BHLHE41
FOS
SIX1
SOX8
MEIS1
E2F6
SOX13
DBX2
RFX4
TCF4
REST
NFIX
POU3F2
MYBL2
MYBL1
E2F8
TFDP1
EZH2
DN − 1
DN − 2
Excitatory
Inhibitory
Neurons
Astrocytes − Early
Astrocyte
Endothelial
Neural-Endothelial
Ependymal
Fibroblast
OPC
RG
Stem
B
FOXA1
0.20
0.15
0.10
0.05
C
FOXA2
0.25
0.20
0.15
0.10
0.05
D
EN1
0.3
0.2
0.1
E
SOX6
0.4
0.3
0.2
0.1
0.0
F
LMX1A
0.5
0.4
0.3
0.2
0.1
0.0

**Supplementary Figure 1**

**Identification of transcription factor activity across cell types in *SNCA* triplication and isogenic control midbrain organoids. (A)** Heatmap of top 5 regulons per cell cluster in SNCA Trip and Iso CTL hMOs determined by pySCENIC analysis. **(B-F)** Expression plots of selected DN transcription factors, highlighting their activity in DN clusters.

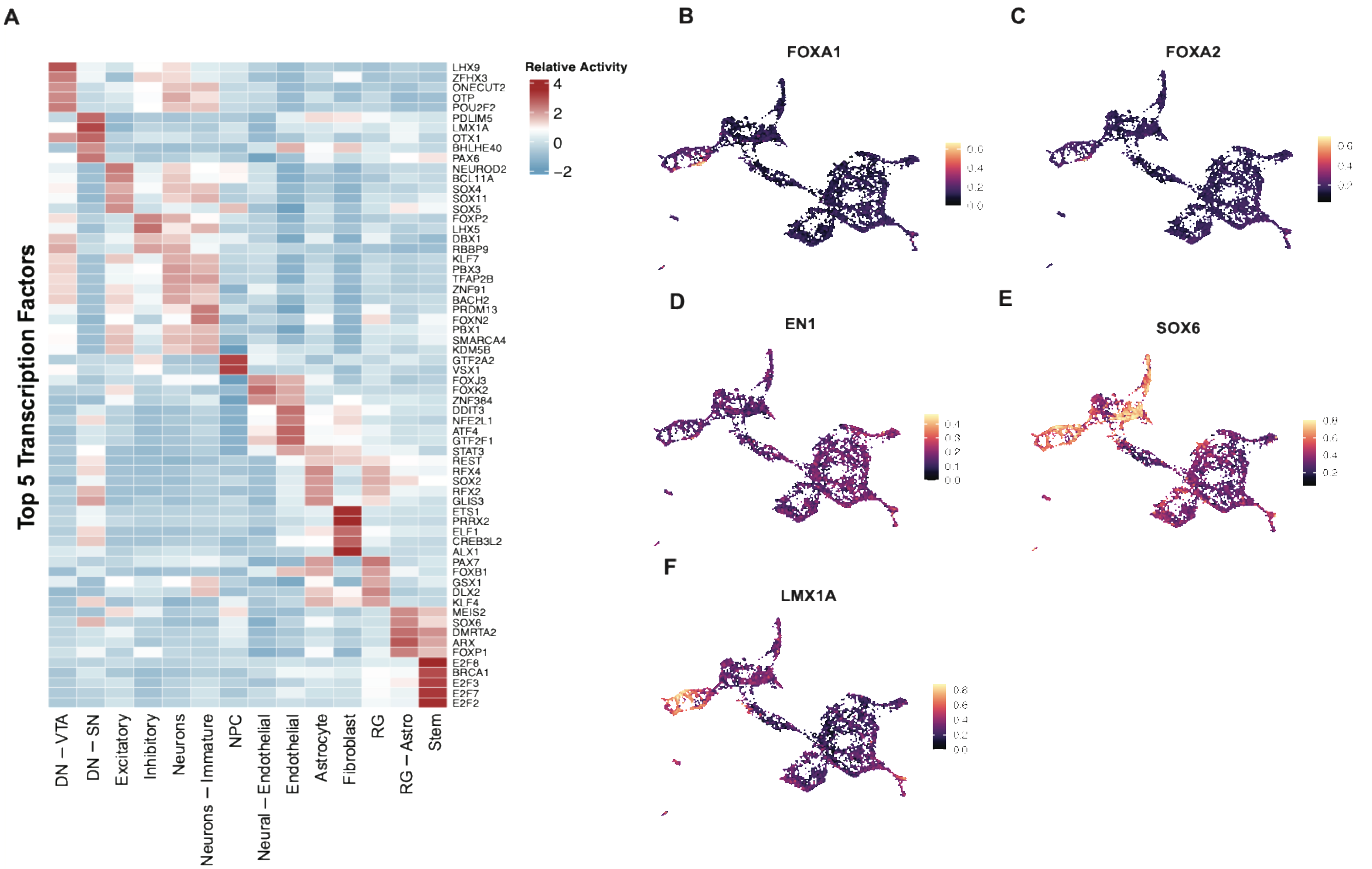

A
Top 5 Transcription Factors
LHX9
ZFHX3
ONECUT2
OTP
POU2F2
PDLIM5
LMX1A
OTX1
BHLHE40
PAX6
NEUROD2
BCL11A
SOX4
SOX11
SOX5
FOXP2
LHX5
DBX1
RBBP9
KLF7
PBX3
TFAP2B
ZNF91
BACH2
PRDM13
FOXN2
PBX1
SMARCA4
KDM5B
GTF2A2
VSX1
FOXJ3
FOXK2
ZNF384
DDIT3
NFE2L1
ATF4
GTF2F1
STAT3
REST
RFX4
SOX2
RFX2
GLIS3
ETS1
PRRX2
ELF1
CREB3L2
ALX1
PAX7
FOXB1
GSX1
DLX2
KLF4
MEIS2
SOX6
DMRTA2
ARX
FOXP1
E2F8
BRCA1
E2F3
E2F7
E2F2
DN – VTA
DN – SN
Excitatory
Inhibitory
Neurons
Neurons – Immature
NPC
Neural – Endothelial
Endothelial
Astrocyte
Fibroblast
RG
RG – Astro
Stem
Relative Activity
4
2
0
−2
B
FOXA1
C
FOXA2
D
EN1
E
SOX6
F
LMX1A

**Supplementary Figure 2**

**Characterization of transcription factor activity across cell types in sporadic Parkinson's disease and control lines.** **(A)** Heatmap of top 5 regulons per cell cluster in SP-PD and CTL hMOs determined by pySCENIC analysis. **(B-F)** Expression plots of selected DN transcription factors, highlighting their activity in DN clusters.

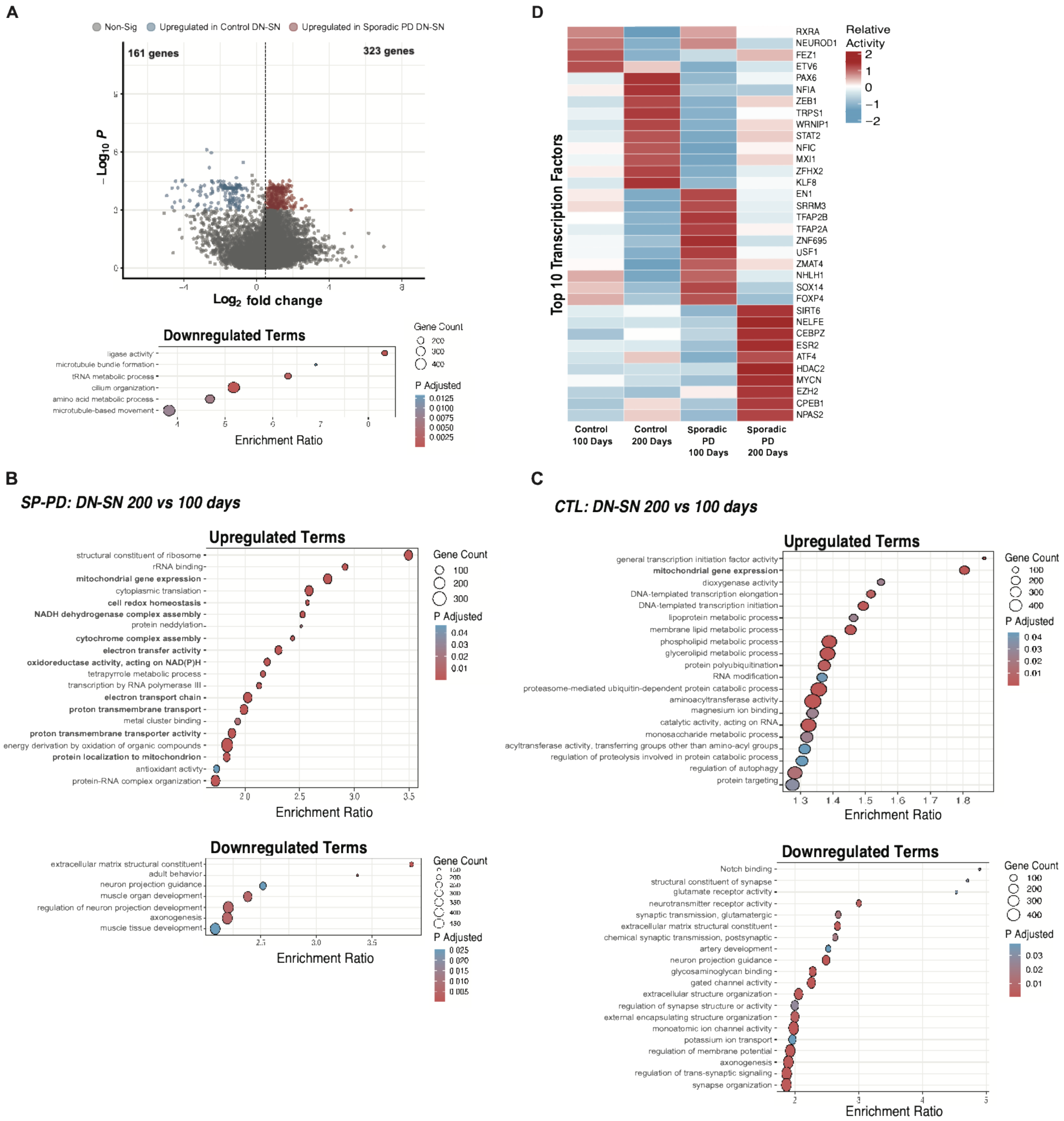
A
Non-Sig
Upregulated in Control DN-SN
Upregulated in Sporadic PD DN-SN
161 genes
323 genes
Log2 fold change
Downregulated Terms
ligase activity
microtubule bundle formation
tRNA metabolic process
cilium organization
amino acid metabolic process
microtubule-based movement
Enrichment Ratio
Gene Count
P Adjusted
D
Top 10 Transcription Factors
Relative Activity
RXRA
NEUROD1
FEZ1
ETV6
PAX6
NFIA
ZEB1
TRPS1
WRNIP1
STAT2
NFIC
MXI1
ZFHX2
KLF8
EN1
SRRM3
TFAP2B
TFAP2A
ZNF695
USF1
ZMAT4
NHLH1
SOX14
FOXP4
SIRT6
NELFE
CEBPZ
ESR2
ATF4
HDAC2
MYCN
EZH2
CPEB1
NPAS2
Control 100 Days
Control 200 Days
Sporadic PD 100 Days
Sporadic PD 200 Days
B
SP-PD: DN-SN 200 vs 100 days
Upregulated Terms
structural constituent of ribosome
rRNA binding
mitochondrial gene expression
cytoplasmic translation
cell redox homeostasis
NADH dehydrogenase complex assembly
protein neddylation
cytochrome complex assembly
electron transfer activity
oxidoreductase activity, acting on NAD(P)H
tetrapyrrole metabolic process
transcription by RNA polymerase III
electron transport chain
proton transmembrane transport
metal cluster binding
proton transmembrane transporter activity
energy derivation by oxidation of organic compounds
protein localization to mitochondrion
antioxidant activity
protein-RNA complex organization
Enrichment Ratio
Gene Count
P Adjusted
Downregulated Terms
extracellular matrix structural constituent
adult behavior
neuron projection guidance
muscle organ development
regulation of neuron projection development
axonogenesis
muscle tissue development
Enrichment Ratio
C
CTL: DN-SN 200 vs 100 days
Upregulated Terms
general transcription initiation factor activity
mitochondrial gene expression
dioxygenase activity
DNA-templated transcription elongation
DNA-templated transcription initiation
lipoprotein metabolic process
membrane lipid metabolic process
phospholipid metabolic process
glycerolipid metabolic process
protein polyubiquitination
RNA modification
proteasome-mediated ubiquitin-dependent protein catabolic process
aminoacyltransferase activity
magnesium ion binding
catalytic activity, acting on RNA
monosaccharide metabolic process
acyltransferase activity, transferring groups other than amino-acyl groups
regulation of proteolysis involved in protein catabolic process
regulation of autophagy
protein targeting
Enrichment Ratio
Downregulated Terms
Notch binding
structural constituent of synapse
glutamate receptor activity
neurotransmitter receptor activity
synaptic transmission, glutamatergic
extracellular matrix structural constituent
chemical synaptic transmission, postsynaptic
artery development
neuron projection guidance
glycosaminoglycan binding
gated channel activity
extracellular structure organization
regulation of synapse structure or activity
external encapsulating structure organization
monoatomic ion channel activity
potassium ion transport
regulation of membrane potential
axonogenesis
regulation of trans-synaptic signaling
synapse organization
Enrichment Ratio
Gene Count
P Adjusted

**Supplementary Figure 3**

**Differential gene expression analysis between sporadic Parkinson's and control midbrain organoids reveals increased mitochondrial gene expression over time and across multiple cell types in Parkinson's lines. (A)** Ontop: Volcano plot highlighting differentially expressed genes between SP-PD and CTL DN-SNs at 100 days. Significance thresholds are PAdj < 0.001 and Log2 fold change >0.5 or <-0.5 Below: Gene ontology analysis shows only pathways associated with downregulated genes. **(B)** Gene ontology analysis between DN-SNs in 100- and 200- day SP-PD hMOs highlighting significantly up- and down- regulated terms. **(C)** Gene ontology analysis between DN-SNs in 100- and 200- day CTL hMOs highlighting significantly up- and down- regulated terms. **(D)** Top differentially active transcription factors in SP-PD and CTL DN-SN at 100 days and 200 days.

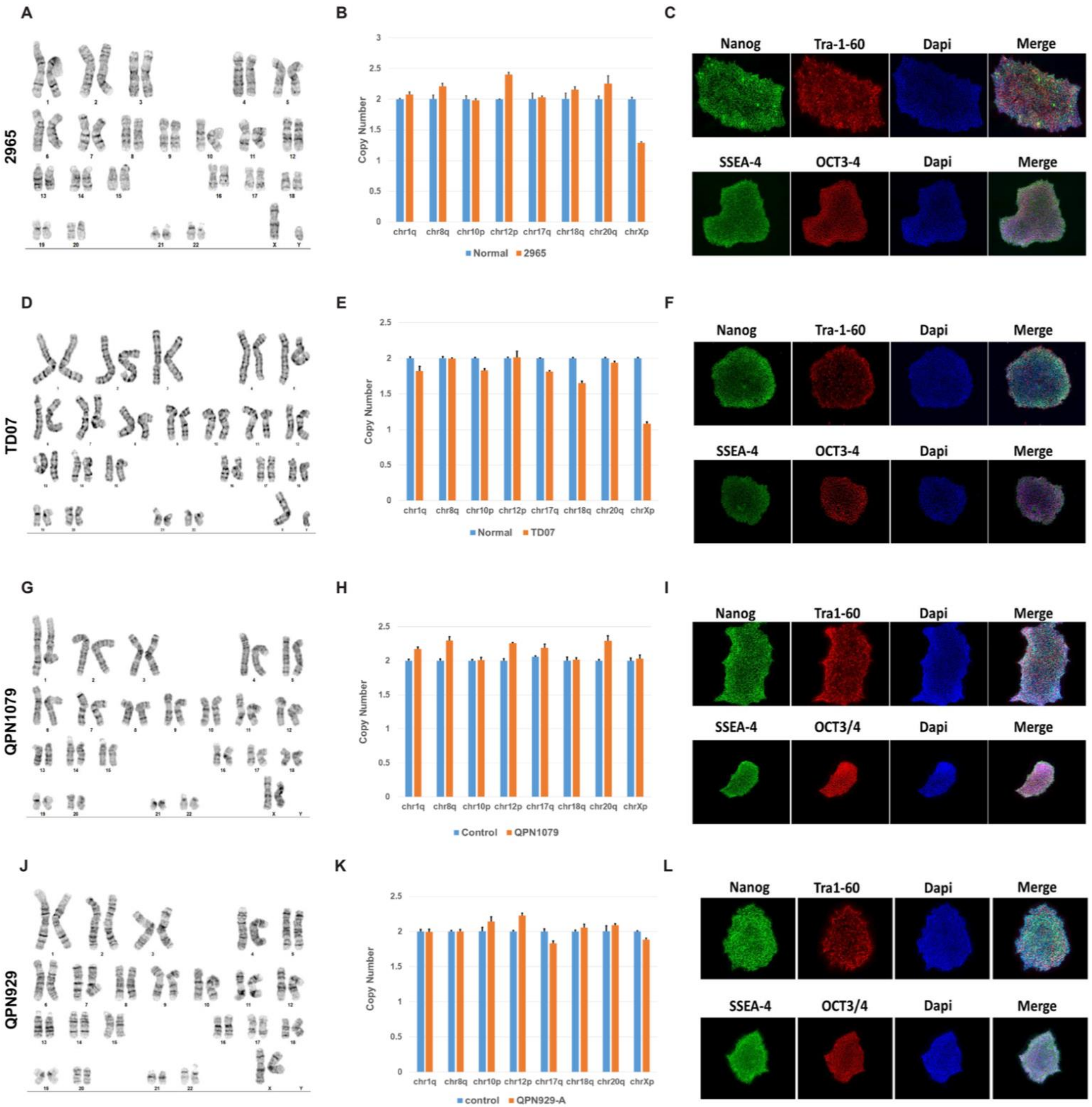

A
2965
B
Copy Number
chr1q
chr8q
chr10p
chr12p
chr17q
chr18q
chr20q
chrXp
Normal
2965
C
Nanog
Tra-1-60
Dapi
Merge
SSEA-4
OCT3-4
Dapi
Merge
D
TD07
E
Copy Number
chr1q
chr8q
chr10p
chr12p
chr17q
chr18q
chr20q
chrXp
Normal
TD07
F
Nanog
Tra-1-60
Dapi
Merge
SSEA-4
OCT3-4
Dapi
Merge
G
QPN1079
H
Copy Number
chr1q
chr8q
chr10p
chr12p
chr17q
chr18q
chr20q
chrXp
Control
QPN1079
I
Nanog
Tra1-60
Dapi
Merge
SSEA-4
OCT3/4
Dapi
Merge
J
QPN929
K
Copy Number
chr1q
chr8q
chr10p
chr12p
chr17q
chr18q
chr20q
chrXp
control
QPN929-A
L
Nanog
Tra1-60
Dapi
Merge
SSEA-4
OCT3/4
Dapi
Merge

**Supplementary Figure 4**

**Quality control of unpublished iPSC lines. (A)** Karyotyping performed on extracted DNA from 2965 (Sporadic PD patient) iPSC line. **(B)** Genomic stability assay performed on extracted DNA from 2965 iPSCs showing normal copy numbers in frequently mutated sites. **(C)** Immunofluorescent staining against pluripotency markers NANOG, TRA1-60, SSEA-4, OCT3-4 in 2965 iPSCs. **(D-F)** Karyotyping, genomic stability assay, and immunofluorescent staining against pluripotency markers in TD07 iPSCs (Sporadic PD patient). **(G-I)** Karyotyping, genomic stability assay, and immunofluorescent staining against pluripotency markers in QPN1079 iPSCs (Sporadic PD patient). **(J-L)** Karyotyping, genomic stability assay, and immunofluorescent staining against pluripotency markers in QPN929 iPSCs (Healthy control).

**Supplementary Table 1**

| Marker | References |
|---|---|
| CALB1 | Luppi et.al 2021; Poulin et.al 2014; Chung et.al 2005; Greene et.al 2005; Fiorenzano et.al 2021; Poulin et.al 2018 |
| CALB2 | Poulin et.al 2014; Chung et.al 2005; Fiorenzano et.al 2021 |
| ADCYAP1 | Poulin et.al 2014; Chung et.al 2005; Greene et.al 2005 |
| BDNF | Greene et.al 2005 |
| CNR1 | La Manno et.al 2016 |
| NEFM | Poulin et.al 2014 |
| NEUROD6 | Chung et.al 2005; La Manno et.al 2016; Poulin et. al 2014 |
| SLC17A6 | Luppi et.al 2021; Poulin et.al 2014; Grimm et.al. 2004; Chung et.al 2005; Tiklova et.al 2019; Salmani et.al. 2024; Poulin et.al 2018 |
| GAD2 | Salmani et.al 2024 |
| GAD1 | Chung et.al 2005; Greene et. al 2005; Tiklova et.al 2019 |
| CBLB | Tang et.al 2023 |
| KCNH1 | Tang et.al 2023 |
| NGB | Luppi et.al 2021 |
| PARM1 | Grimm et.al 2004 |
| PBX3 | Kamath et.al 2022 |
| POU3F1 | Kamath et.al 2022; La Manno et.al 2016 |
| SORCS3 | Kamath et.al 2022; Tang et.al 2023 |
| TGFB1 | Kamath et.al 2022 |
| TMEFF2 | Kamath et.al 2022; Tang et.al 2023 |
| SOX6 | Luppi et.al 2021; Poulin et.al 2014; Grimm et.al 2004; Chung et.al 2005; La Manno et.al 2016; Salmani et.al 2024; Poulin et.al 2018 |
| ALDH1A1 | Chung et.al 2005; La Manno et.al 2016; Salmani et.al 2024; Tiklova et.al 2019; Poulin et.al 2014; Poulin et.al 2018 |
| NDNF | La Manno et.al 2016; Salmani et.al 2024; Poulin et.al 2018 |
| SERPINE2 | Luppi et.al 2021; Hook et.al 2018 |
| ANXA1 | La Manno et.al 2016; Fiorenzano et.al 2021; Salmani et.al 2024; Poulin et.al 2018 |
| CADPS2 | Tang et.al 2023; Smajic et.al 2022 |
| COL11A1 | Tang et.al 2023 |
| CTSD | Greene et.al 2005 |
| ENO1 | Greene et.al 2005 |
| ENO3 | Greene et.al 2005 |
| FGF20 | Luppi et.al 2021 |
| LRRK2 | Tang et.al 2023 |
| PSD3 | Tang et.al 2023 |
| RYR3 | Tang et.al 2023 |
| SH3GL2 | Tang et.al 2023 |

**Supplementary Table 2**

**Description of iPSC cell lines**

| Line Name | Description | Source | Sex and age | Quality Control |
|---|---|---|---|---|
| AST23 | SNCA Triplication (Patient line) | Dr. T. Kunaths laboratory, The University of Edinburgh | Female 50-54 years old | Mohamed et.al. 2021[27] Devine et.al. 2011[75] |
| AST23-2KO | Wild-type (CRISPR-corrected Isogenic) | Dr. T. Kunaths laboratory, The University of Edinburgh | Female 50-54 years old | Mohamed et.al. 2021[27] Chen et.al. 2018[76] |
| AIW001 | Control | Early Drug Discovery Unit, McGill University | Female 48 years old | Chen et.al. 2021[80] |
| AIW002 | Control | Early Drug Discovery Unit, McGill University | Male 37 years old | Chen et.al. 2021[80] |
| QPN929 | Control | Early Drug Discovery Unit, McGill University | Female 50 years old | Supplementary Fig.4J-L |
| TD22 | Control | Early Drug Discovery Unit, McGill University | Male 59 years old | Chen et.al. 2021[80] |
| 2965 | Sporadic PD | Early Drug Discovery Unit, McGill University | Male 67 years old | Supplementary Fig.4A-C |
| QPN1079 | Sporadic PD | Early Drug Discovery Unit, McGill University | Female 72 years old | Supplementary Fig.4G-I |
| TD07 | Sporadic PD | Early Drug Discovery Unit, McGill University | Male 70 years old | Supplementary Fig.4D-F |

**Supplementary Table 3**

**Single cell reference datasets**

| Reference | Description | Data Type | Cell Types |
|---|---|---|---|
| Kamath et.al 2022 *Nat Neuro.* | Human postmortem brain tissue | snRNAseq | Enriched dopamine neurons |

| | | | |
|---|---|---|---|
| Jerber et.al 2021 *Nat Gen.* | iPSC derived dopamine neurons | scRNAseq | Dopamine neurons |
| Nowakowski et.al 2017 *Science* | Human fetal cortical tissue | scRNAseq | Broad cortical cell types |
| LaManno et.al 2016 *Cell* | Human fetal ventral midbrain tissue | scRNAseq | Broad midbrain cell types |
| Xu et.al 2025 *Nat Gen.* | Human organoid atlas | scRNAseq | Broad brain cell types |

**Supplementary Table 4**

**Cell type marker list**

| **Cell type** | **Marker list** |
|---|---|
| Dopaminergic | TH, SLC18A2, FOXA2, EN1, DDC, PITX3, LMX1A, LMX1B, NR4A2 |
| Excitatory | SLC17A6, SLC17A7, NEUROD2, CALB2, GRID1 |
| Inhibitory | GAD1, GAD2, GRIK1, SLC32A1, SLC6A1, PRL, SOX14 |
| Neurons | STMN2, SYP, HOMER2, RBFOX3 |
| Astrocytes | GFAP, AQP4, S100B, AMIGO2, EGFR, SOX9, ALDH1L1 |
| Endothelial | ICAM1, ALPL, ENG, FLT1, GATA4, GATA6, FOXF1 |
| Ependymal | CCDC78, CXCL5, CAPN8, FOXJ1, PIFO, DYNLRB2 |
| Fibroblast | COL15A1, COL3A1, COL1A2, COL6A3, MGP, MMP2, MMP19, CCN4 |
| Oligodendrocyte | OLIG1, OLIG2, NG2, MBP, PLP1, GALR1 |
| Radial Glia | SLC1A3, PAX6, GLI3, HES5, NTN1, LRRC10B, RPE65, GMPR, PAX3 |
| Stem | MKI67, NANOG, PCNA |
| Progenitor | NES, HES1, SOX2 |

**Supplementary Table 5**

**Summary of antibodies used**

| **Antibody** | **Company** | **Cat Number** | **Application** | **Concentration** |
|---|---|---|---|---|
| TH anti-Ch | Abcam | Ab76442 | IF | 1:500 |

| | | | | |
|---|---|---|---|---|
| ALDH1A1 anti-Rb | Sigma | HPA002123 | IF | 1:500 |
| CALB1 anti-GP | Synaptic Systems | 214 004 | IF | 1:1000 |
